# Explore or reset? Pupil diameter transiently increases in self-chosen switches between cognitive labor and leisure in either direction

**DOI:** 10.1101/379214

**Authors:** Johannes Algermissen, Erik Bijleveld, Nils B. Jostmann, Rob W. Holland

## Abstract

When people invest effort in cognitive work, they often keep an eye open for rewarding alternative activities. Previous research suggests that the norepinephrine (NE) system regulates such trade-offs between exploitation of the current task and exploration of alternative possibilities. Here we examine the possibility that the NE-system is involved in another trade-off, i.e., the trade-off between cognitive labor and leisure. We conducted two pre-registered studies (total *N* = 62) in which participants freely chose to perform either a paid 2-back task (labor) vs. a non-paid task (leisure), while we tracked their pupil diameter—which is an indicator of the state of the NE system. In both studies, consistent with prior work, we found (a) increases in pupil baseline and (b) decreases in pupil dilation when participants switched from labor to leisure. Unexpectedly, we found the same pattern when participants switched from leisure back to labor. Both increases in pupil baseline and decreases in pupil dilation were short-lived. Collectively, these results are more consistent with a role of norepinephrine in reorienting attention and task switching, as suggested by network reset theory, than with a role in motivation, as suggested by adaptive gain theory.

---

In their pursuit of rewards, such as food, all organisms are continuously confronted with two choice options: they can either keep exploiting their current location to harvest rewards, or they can leave their current place to explore the environment for potentially more attractive alternatives. These two strategies, called exploitation and exploration, can both contribute to maximizing rewards in the long-term: When organisms *exploit*, they aim to maximize rewards by sticking with the activity that they believe yields the highest payoff at the moment; when organisms *explore*, they aim to maximize rewards by gathering information on whether other activities yield higher returns. This fundamental dilemma between *exploitation* and *exploration* plays a central role in foraging models, which have a rich history in behavioral ecology (Charnov, 1976). Foraging models have proven valuable in various fields, including artificial intelligence, psychology, and neuroscience (Calhoun & Hayden, 2015; Hayden, 2018; Hills, Todd, Lazer, Redish, & Couzin, 2015), and have recently been suggested as a general framework for understanding value-based choice (Hayden, 2018; Hunt & Hayden, 2017; Rushworth, Kolling, Sallet, & Mars, 2012).

A limitation of previous foraging experiments with humans is that these experiments did not account for the *effort costs* of different activities (Constantino & Daw, 2015; Kolling, Behrens, Mars, & Rushworth, 2012). In real life, obtaining higher rewards often requires higher effort exertion (e.g., longer, faster, or more intense performance in sports), and thus higher effort is often strategically invested for higher reward prospects (Bijleveld, Custers, & Aarts, 2009, 2012). Similar to reward pursuit in foraging, also physical (Meyniel, Sergent, Rigoux, Daunizeau, & Pessiglione, 2013) and mental (Kool & Botvinick, 2014) effort mobilization are not constant over time. Instead, effort mobilization is interrupted by breaks that might serve the recreation from effort expansion (Jett & George, 2003). It thus makes sense to conceptualize decisions for rewards not just as a trade-off between *exploration* and *exploitation*, but also as a trade-off between *cognitive labor* (i.e., exerting effort to attain a reward) and *cognitive leisure* (i.e., carrying out a non-demanding and non-profitable activity, such as relaxing; Kool & Botvinick, 2014, 2018). Note that labor and leisure are conceptualized in relative terms, such that a task is a leisure task if it provides relief from effort mobilization in another (labor) task; no task is intrinsically labor or leisure.

It is important to note that these two dilemmas—exploration vs. exploitation and labor vs. leisure—are conceptually independent. After all, exploitation vs. exploration concerns the question of how to maximize reward, whereas labor vs. leisure concerns the question of how to optimally allocate effort. Put differently, the exploration–exploitation dilemma emerges when people face a choice between (a form of) short-term reward pursuit and (a form of) information gathering, whereas the labor–leisure dilemma emerges when people face a choice between some high-effort activity and some activity that provides relief. Because of their independence, the exploitation–exploitation and labor–leisure trade-offs do not directly map onto each other. That is, labor activities can sometimes be exploitative (e.g., when working on a spreadsheet helps to make progress to a work-related goal, and thus, is continued), but at other times exploratory (e.g., when working on a spreadsheet is chosen as a potentially rewarding distraction from some other activity). Similarly, leisure activities can be exploitative (e.g., when talking to colleagues at work is rewarding, and thus, is continued) but also exploratory (e.g., when talking to colleagues is chosen as a potentially rewarding distraction from some other activity).

In this research, we examine the particular case in which people are faced with the choice between high-effort cognitive work that yields a higher expected reward (i.e., labor) and taking a low-effort break that yields a lower expected reward (leisure). In our view, this case is prototypical of many real-life decisions under uncertainty. In such cases, labor may often co-occur with exploitation (since labor tasks reliably yield external rewards), and leisure may often co-ccur with exploration (since leisure tasks yield no external reward, but provide opportunities to detect other, more intrinsic rewards, e.g., encouraging social interactions). Our research concerns the particular cases in which this mapping holds, and is agnostic about other cases in which this mapping is reversed or more dynamic (e.g., when this mapping changes over time because the rewards associated with labor and leisure change over time). By studying these particular cases, we examine a potential connection between the two trade-offs. In particular, using a labor-leisure paradigm, we investigate whether the neural mechanisms that people seem to use to balance exploitation and exploration (i.e., the locus-coeruleus norepinephrine system) may also be used for trading off cognitive labor and leisure.

In our research, we go beyond prior work in two ways. First, we used a dual-task paradigm with two qualitatively different tasks. We chose to use this paradigm to model decisions in real life, where labor (e.g., working on a spreadsheet) usually comprises a vastly different activity than leisure (e.g., talking to colleagues). Second, we examined transitions between labor and leisure in both directions. That is, we not only examined decisions to take breaks (while working), but also examined when people decided to start working again (after taking a break). An important feature of our studies is that analysis plans were pre-registered before data collection. In what follows, we introduce the major biological theory linking the exploration-exploitation trade-off to neural processes, describe previous human research supporting this theory, and explain why our paradigm allows for a more comprehensive investigation of NE levels when humans switch between labor and leisure.

A major theory linking the trade-off between exploration and exploitation to neural mechanisms is *adaptive gain theory*, which connects these behavioral states to qualitatively different neural states of the locus coeruleus–norepinephrine (LC-NE) system (Aston-Jones & Cohen, 2005; Cohen, McClure, & Yu, 2007). According to adaptive gain theory, *exploitation* is driven by a *phasic* mode of locus coeruleus (LC) activity. In the phasic mode, baseline norepinephrine (NE) levels in the LC are moderate, but bursts of NE release occur in response to task-related stimuli. Given that NE release in cortical areas increases neural responsivity to incoming information (Berridge & Waterhouse, 2003; Servan-Schreiber, Printz, & Cohen, 1990), this pattern is likely adaptive: it helps animals to process task-relevant information. In contrast, *exploration* is driven by a *tonic* mode of LC activity. In the tonic mode, baseline NE levels are chronically elevated, and bursts of NE (in response to task-relevant stimuli) are attenuated or even absent. It has been suggested that this pattern may widen attention, allowing people to better detect task-irrelevant—but potentially rewarding—stimuli (Cohen et al., 2007).

Importantly, NE levels are correlated with pupil diameter in both monkeys (Joshi, Li, Kalwani, & Gold, 2016; Rajkowski, Kubiak, & Aston-Jones, 1994) and humans (Murphy, O’Connell, O’Sullivan, Robertson, & Balsters, 2014; Murphy, Robertson, Balsters, & O’Connell, 2011). For example, in line with predictions from adaptive gain theory, previous research found that pupil diameter covaried with task engagement and disengagement in an auditory discrimination task (Gilzenrat, Nieuwenhuis, Jepma, & Cohen, 2010). In this task, participants judged which of two tones had a higher pitch. Task difficulty continuously increased while rewards decreased with every error a participant made. At any time, participants could reset task settings by pressing an “escape” button, which the authors interpreted as exploration behavior. Changes in pupil dilation were also observed in transitions between bandit gambling machines (Jepma & Nieuwenhuis, 2011) and in a task that required people to solve Raven’s Matrices (Hayes & Petrov, 2016). Overall, in these studies, *exploitation* was characterized by relatively low baseline pupil diameter, combined with large pupil dilations in response to task stimuli. By contrast, *exploration* was characterized by high baselines and smaller dilations in response to task-relevant stimuli.

While the studies described in the previous paragraph link the LC-NE system, and particularly pupil diameter as an observable correlate, to exploration–exploitation dilemmas, it is not yet clear whether the LC-NE system is involved in people’s decisions to switch between labor and leisure. After all, prior studies operationalized exploration as brief transitions between extended phases of exploitation, making it impossible to disentangle disengagement from the previous task vs. re-engagement in the next task. Also, these studies manipulated task payoff and difficulty, thus incentivizing all participants to start exploration at a defined moment, instead of keeping the environment constant and tracking individual differences in participants’ natural drive for exploration. In contrast, we studied participants’ self-directed decisions to take a break (vs. to continue working), which are arguably similar in structure as exploration–exploitation dilemmas, because agents have to decide whether to (a) stay with a current activity and its payoff or (b) to quit this activity and explore the environment for more rewarding alternatives—at the risk of wasting time and foregoing rewards. Indeed, in research on fatigue and effort, the LC-NE system has been mentioned as a candidate mechanism for understanding labor-leisure transitions (Inzlicht, Schmeichel, & Macrae, 2014; Kurzban, Duckworth, Kable, & Myers, 2013). Yet, to our knowledge, this possibility has not been tested.

## The present research

Based on adaptive gain theory, we hypothesized that labor-to-leisure transitions (i.e., decisions to take a break from work) are preceded by (a) increases in pupil baseline and (b) decreases in pupil dilation. For leisure-to-labor transitions (i.e., decisions to start working after taking a break), predictions are somewhat less straightforward. On the one hand, based on adaptive gain theory, we would predict that baselines stay high during the leisure phase to ensure a broadened attention that facilitates the detection of alternative activities. However, when switching back to labor, people need to focus on the labor task only, so that (a) pupil baseline should decrease and (b) pupil dilations should increase again. On the other hand, one could also pose that leisure-to-labor transitions are not different from labor-to-leisure transitions in that they constitute a case of task-switching. A role of NE in task switching has been suggested by *network reset theory* (Bouret & Sara, 2005; Sara & Bouret, 2012), which suggests that NE is primarily released when people detect unexpected changes in the environment. When these changes happen, NE promotes reorientation towards stimuli that have become relevant in the new task environment, facilitating adaptation to new demands. So far, it has not been explored whether NE could have a similar function in humans’ self-directed decisions to switch tasks. If this was the case, we would expect (a) increases in pupil baseline and (b) decreases in pupil dilation preceding both labor-to-leisure and leisure-to-labor transitions, because network reset theory assumes NE processes to be independent of the activity performed previously and the activity to-be-performed after the switch. By investigating leisure-to-labor transitions, our research directly tested adaptive gain theory and network reset theory against each other.

We operationalized *labor* as a 2-back memory task and *leisure* as an attractiveness rating task, following a paradigm by Kool and Botvinick (2014). We employed both linear and additive mixed models, which allowed us to more closely examine the onset and duration of pupil shifts. To shed light on functional role of the processes underlying pupil changes, we conducted exploratory analyses in which we correlated the magnitude of pupil changes with individual differences in self-reported procrastination and action orientation in everyday life. To this end, we used well-validated questionnaires (Kuhl, 1994; Kuhl & Fuhrmann, 1998; Steel, 2010). We chose personality dimensions that reflected how readily people engaged into effortful tasks, and how they dealt with fatigue following from effort. We explored whether people who successfully dealt with these challenges in everyday life would show a different pupil pattern around switches than people who had problems with initiating or maintaining effort mobilization. These analyses were exploratory and served hypothesis-generation for future studies.

Below, we report Methods and Results for Studies 1 and 2 together, as protocols in both studies were identical except for two brief control assessments administered following the main task in Study 2. In our pre-registration for Study 1,^1^ based on previous literature (Gilzenrat et al., 2010), we expected pupil shifts to occur prior to task switches, which would have allowed us to predict switches prospectively based on pupil diameter. As our hypotheses found no evidence in the pre-registered time window, we explored them in a later time window centered around the behavioral switch. We pre-registered these analyses for Study 2^2^ to replicate our exploratory findings from Study 1.

## Methods

### Participants

In Study 1, 35 participants completed a 50-minute study in exchange for a €7.50 voucher and an extra cash payment of up to €5, depending on their task performance. We recruited participants in the age range 17–30 who had normal or corrected-to-normal vision (using contact lenses), understood English, and did not suffer from neurological disorders. In line with pre-registered exclusion criteria (see Supplementary Online Material S1), four participants were excluded from analyses, so that the final sample consisted of 31 participants (68% female, *M_age_* = 22, *SD_age_* = 2.7).

In Study 2, 35 participants completed a 60-minute study in exchange for a €10 shopping voucher and an extra cash payment of up to €5. Participants were recruited from the same population as in Study 1, using the same exclusion criteria (see S2). Four participants were excluded from analyses, so that the final sample consisted of 31 participants (65% female, *M_age_* = 22, *SD_age_* = 3.1). Studies were approved by the local ethics review board.

### Procedure

Participants were welcomed in a room equipped with a stationary SMI iView X infrared eye-tracker (SensoMotoric Instruments, Teltow, Germany) sampling at 500 Hz. We placed this device’s chin rest 72 cm away from a 24-inch monitor, on which the task stimuli were presented (using a script programmed in PsychoPy; Peirce, 2007). After they were seated, participants first completed a 9-point calibration of the eye-tracking device, followed by a 10 min practice phase, in which participants were familiarized with the labor and leisure tasks (which were first practiced separately), but also with labor-to-leisure and leisure-to-labor switches.

Next, participants completed the test phase, which took 25 min and consisted of 500 trials. During both the practice and test phases, pupil size of their dominant eye was tracked. In Study 2, additionally, participants completed two control assessments of 80 trials each to exclude alternative interpretations of the findings of Study 1. Afterwards, in both studies, participants rated both tasks on the dimensions *work* and *fun* using a 7-point scale. We assumed that the *work* ratings reflected whether participants perceived the labor task as more effortful than the leisure task, which would mean that our task successfully fulfilled the criterion for a labor-leisure trade-off. In contrast, the *fun* ratings assessed participants’ intrinsic motivation (opposed to the extrinsic monetary incentives) to do a task. These fun ratings were important to exclude the possibility that participants performed the labor task not because it returned monetary rewards, but because it was more fun than the leisure task. If participants were motivated by fun instead of maximizing rewards, neural mechanisms different from those designed to deal with exploitation-exploration dilemmas might have been involved. Afterwards, participants filled in questionnaires assessing their procrastination tendency and action vs. state orientation. Finally, they were debriefed and paid.

### Labor-Leisure Task

During the test phase, in each trial, participants could choose between performing an effortful 2-back task (labor task) or an easy attractiveness-rating task (leisure task; adapted from Kool & Botvinick, 2014). For each trial they spent on the labor task, they received 1 cent extra cash payment. It was announced in advance that participants with accuracy levels below 75% in the labor task would receive no extra payment, but that previous participants were able to easily achieve this level if they tried hard. This threshold was implemented in order to discourage participants from resting during the labor task and encourage switching to the leisure task in case they felt like taking a break. Participants’ responses during the leisure task did not affect their monetary payout, allowing for an actual break from paid work.

Participants selected and performed tasks using a joystick. Specifically, participants selected tasks by moving the joystick sideways. When the joystick was moved to the left (or right; counterbalanced), participants performed the labor task; to the right (or left; counterbalanced), participants performed the leisure task. Both tasks were presented as labels at the upper corners of the screen (“2-back” and “attractiveness”). As a reminder, the currently-selected task was surrounded by a white frame. It was only possible to switch tasks when no stimulus was displayed, in order to prevent participants from employing the strategy of first viewing a face and then choosing which task to perform. Participants indicated responses via the single backward trigger of the joystick. We chose to use a joystick in line with the original paradigm by Kool and Botvinick (2014). Compared to a keyboard, a joystick reduces the chance that participants accidently press a different key than they intend; after all, when using a stationary eye tracking device with a chin and head rest, participants could not visually check their manual responses.

Regardless of the position of the joystick, on each trial, participants saw the following stimuli: a fixation cross (200 ms), a mask stimulus (800 ms), a face (1500 ms), and a blank screen (400–600 ms). Figure 1 depicts the course of one trial. When participants had selected the labor task, they were required to press the joystick trigger when the current face was the same face as two faces before (i.e., when the current face was a 2-back target; in case of a nontarget, they were required to refrain from responding). When participants had selected the leisure task, they were asked to press the joystick trigger when they saw an attractive face.

**Figure 1.**
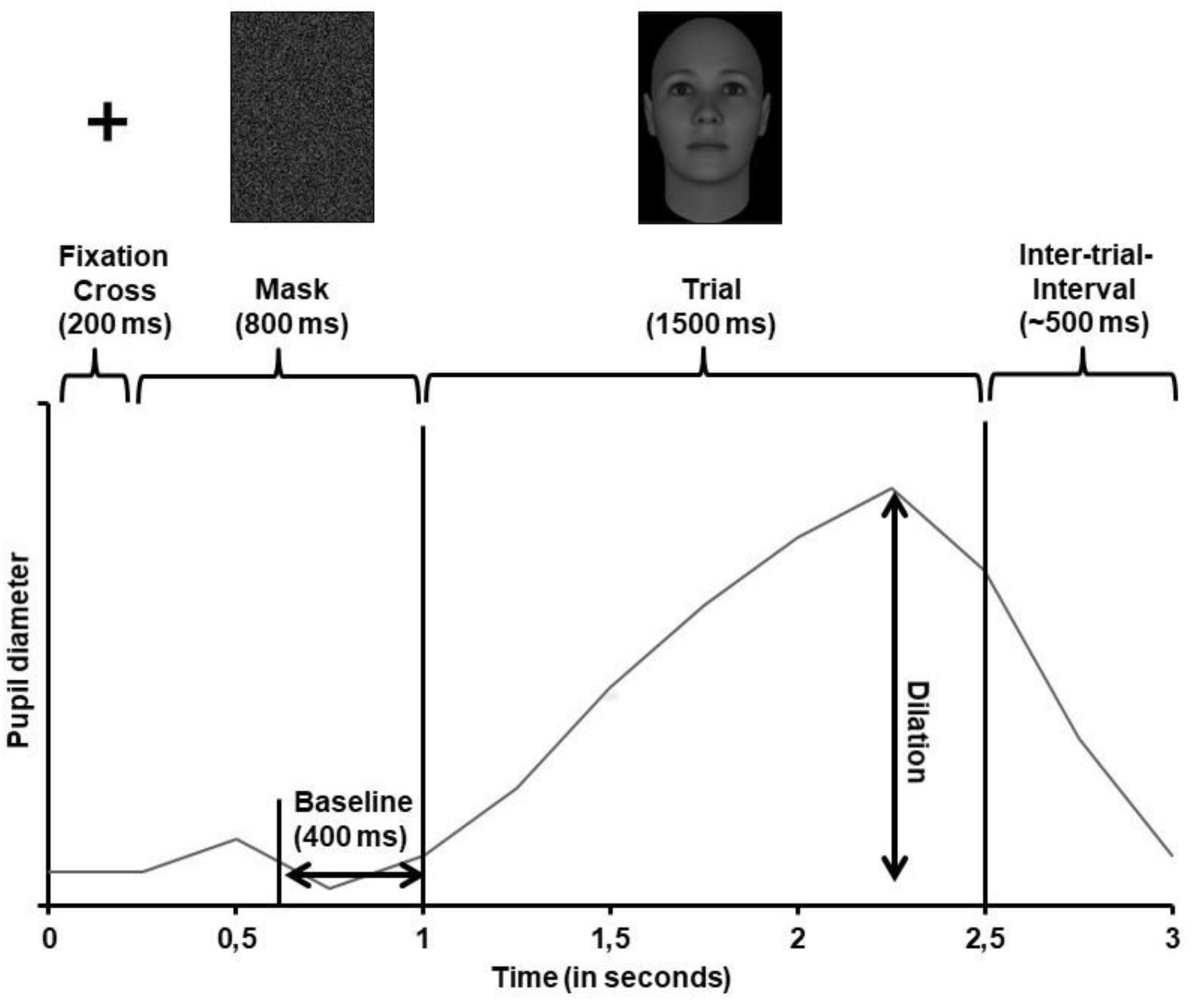
Overview of the time course of one trial in the labor-leisure task, including the parts of the trial in which we measured baselines and dilations, respectively. Face stimulus taken from Said and Todorov (2011).

### Stimuli

Faces were selected from the Attractiveness Model Database (Said & Todorov, 2011), which contains artificially generated faces. Of the 20 faces used, 10 were drawn from the 5% most attractive faces of the database; the other 10 from the 5% least attractive faces. Ten faces were male faces; the other 10 were female faces. All stimuli were matched on mean luminance and had the same size (400 x 400 pixels). The mask consisted of the pixels of one of the faces randomly rearranged, resulting in the same mean luminance. For all practice phases and the final test phase, we generated a pseudo-random order of faces, with a 30% chance of a 2-back trial occurring. Fifty percent of trials showed attractive faces.

### Control Assessments

In Study 2, after the labor-leisure task, participants additionally completed two control assessments of 80 trials (5 min) each. Both conditions were identical to the labor-leisure task in trial structure, stimuli, and task labels, but different in instructions.

First, in the *motor control assessment*, participants were instructed to respond to one particular face (and not to any other face) by moving the joystick in a way as if they switched between tasks in the labor-leisure task. This particular face was displayed on trials 5, 15, … 75, with each of these trials surrounded by 10 trials without any action, allowing for a similar analysis of “switches” as in the labor-leisure task. With this basic vigilance task, we intended to test whether pupil changes were driven by the motor activity occurring when moving the joystick.

Second, in the *visual control assessment*, participants were instructed to passively watch a series of trials without any action. In trials 5, 15, … 75, the frame highlighting the 2-back or attractiveness-rating task label was automatically shifted to the respective other task label. With this control assessment, we intended to test whether watching the moving frame was sufficient to induce the observed pupil changes.

### Questionnaires

In both studies, after people had completed the tasks and left the eye-tracker, we administered three questionnaires:

The Irrational Procrastination Scale (IPS; Steel, 2010) measures general procrastination tendencies using nine items such as “I often regret not getting to task sooner”. People responded on a 7-point scale ranging from strongly disagree (1) to strongly agree (7).

The Action Control Scale (ACS-24; Kuhl, 1994) consists of 24 descriptions of scenarios, with two possible strategies to act in each situation. Participants are asked to select the strategy that describes best how they would react. It comprises two subscales: the action orientation subsequent to failure vs. preoccupation (AOF) subscale contains scenarios about dealing with failures such as “When several things go wrong on the same day”, where the action-oriented response is “I just keep on going as though nothing had happened” and the state-oriented one is “I don’t know how to deal with it.” In contrast, the prospective and decision-related action orientation vs. hesitation (AOD) subscale contains items related to planning and starting activities such as “When I have an obligation to do something that is boring and uninteresting” with the action-oriented response “I do it and get it over with” and the state-oriented response “It usually takes a while before I get around to doing it.”

Furthermore, we selected three subscales from the Self-Government Inventory (SSI-K3; Kuhl & Fuhrmann, 1998): the self-regulation (competence) subscale relates to feelings of autonomy, intrinsic motivation, and dealing with nervousness and contains items such as “When my perseverance subsides, I know exactly how to motivate myself again.” The self-control subscale comprises items about planning, prospection, and self-confidence such as “If I have a lot to do, I work according to a plan (i.e. I have a schedule for my tasks).” The volitional development (action development) subscale consists of items reflecting initiative and readiness to act in contrast to postponement and procrastination such as “If something has to be done, I begin doing it immediately.” Participants rated how much each item applied to themselves on a four-point scale from “not at all” to “completely.” For each of the scales, we calculated participants’ average scores.

### Data Analysis

#### Pre-processing of eye-tracking data

Pre-processing of the pupil raw data included removing values of zero, removing abnormally fast pupil changes, deleting outliers, and finally imputing missing values using linear interpolation. Our exact analysis script was built on previous experimental work (Bijleveld, 2018) and it is available on https://osf.io/b9z4c/ (Study 1) and https://osf.io/ukgsh/ (Study 2).

#### Pupil measures

For each trial, baseline pupil diameter and maximal pupil dilation were computed. Baseline pupil diameter was defined as the average pupil size during the last 400 ms of the mask presentation. Pupil dilation was defined as the difference between the maximal pupil size during face stimulus presentation and the baseline pupil diameter (for an illustration, see Figure 1).

#### Generalized additive mixed models

To account for auto-correlation in the pupil data, we followed recent suggestions (Baayen, Vasishth, Kliegl, & Bates, 2017) to fit generalized additive mixed models (GAMMs) to time-series data using the mgcv package (version 1.8.22; Wood, 2017) in R (version 3.4.3; R Core Team, 2017). The unit of analysis were single trials. Trials were nested in *bouts*, with a bout formed by the five trials before and five trials after a switch. We used either pupil baseline or pupil dilation as the dependent variable, each maximum normalized per person.^3^ We used (a) trial number relative to switch and (b) the interaction between trial number and switch type (labor-to leisure vs. leisure-to-labor) as predictors. We fitted GAMMs with random slopes for trial number for each bout of each participant, which effectively reduced auto-correlation to levels < .15.^4^ When testing for differences between switch types, we coded the predictor switch types as an ordered factor.

Note that GAMMs fit a smooth curve (consisting of thin plate regression splines) through all data points and test whether this curve is significantly different from a straight line (i.e. from zero) at any time point during the selected time window. Thus, GAMMs do not qualify the shape (e.g., linear vs. curvilinear) or direction (e.g., increase vs. decrease) of a change. Hence, we additionally fitted linear mixed-effects models to test for linear increases or decreases in pupil baseline and pupil dilation. In our interpretations, we gave priority to the results of the GAMMs as those (a) account for auto-correlations, and (b) our hypotheses were agnostic about how early and how long the predicted changes would occur.

#### Linear mixed-effects models

We fitted linear mixed-effects models (LMEMs) using the lme4-package (version 1.1.15; Bates, Mächler, Bolker, & Walker, 2015) with pupil baseline or pupil dilation as outcome variable and trial number relative to switch as sole predictor. When comparing the effects of switch types, we added the factor switch type and the interaction between trial number and switch type. Both outcome measures and relative trial numbers were standardized, so that regression coefficients can be interpreted as standardized regression weights. For the factor switch type, we employed sum-to-zero coding. Models contained a maximal random effects structure (Barr, Levy, Scheepers, & Tily, 2013), with random intercepts and random slopes of trial number, switch type, and their interaction, both for each participant and for each bout of adjacent trials of each participant, and with all possible random correlations. We computed type-3 like *p*-values using *F*-tests with Kenward-Roger approximation for degrees of freedom (Singmann, Bolker, Westfall, & Aust, 2017).

#### Power estimation

We checked in a small pilot study (*N* = 7) whether participants followed the task instructions and switched between tasks often enough to yield sufficient power for testing our hypotheses with *N =* 30. Participants switched on average 8.4 times from labor to leisure. Given *N* = 30 and our initial hypothesis of analyzing the last ten but one trial before switches (see below), we expected to obtain 2,268 usable trials. In our pilot data, we obtained intra-class correlations of .67 for baselines and .37 for dilations. Following Aarts, Verhage, Veenvliet, Dolan, and van der Sluis (2014), we estimated effective sample sizes of 357 trials for baselines and 597 trials for dilations, which allowed us to detect effects of β > .14 for baselines and β > .11 for dilations with 80% power (power sensitivity analysis for linear bivariate regression in G*Power 3; Faul, Erdfelder, Lang, & Buchner, 2007).

## Results

### Manipulation Checks

#### Task ratings

In Study 1, participants perceived the 2-back task (*M* = 5.87, *SD* = 0.85) as more work compared to the rating task (*M* = 2.00, *SD* = 1.03), *t*(30) = 17.13, *p* < .001, *d* = 3.08, but also as significantly less fun (*M* = 2.58, *SD* = 0.92) compared to the rating task (*M* = 3.70, *SD* = 1.53), *t*(30) = −4.25, *p* < .001, *d* = −0.76. Also in Study 2, participants perceived the 2-back task as more work (*M* = 5.32, *SD* = 1.45) than the rating task (*M* = 2.32, *SD* = 1.25), *t*(30) = 8.09, *p* < .001, *d* = 1.45, while fun ratings did not significantly differ between the 2- back task (*M* = 2.84, *SD* = 1.37) and the rating task (*M* = 2.94, *SD* = 1.34), *t*(30) = 0.31, *p* = .756, *d* = −0.06. This resulted from a lower “fun”-rating of the rating task compared to Study 1, which might have been due to the fact that in Study 2, between the test phase and the ratings, participants completed the control assessments for another ten minutes. Hence, their memory of the tasks might have faded. Overall, we conclude that in both studies, (a) the task successfully implemented labor in contrast to leisure and (b) the task established a context in which not fun, but reward maximization, drove participants’ choices as to which task to perform.

#### Task performance

In Study 1, on average, participants spent 396 (*SD* = 53) out of 500 trials on the labor task; they switched 7.97 times (*SD* = 7.64) from labor to leisure. In the labor task, participants gave on average 84% (*SD* = 4%) correct responses. In the leisure task, they responded in 27% (*SD* = 14%) of trials, indicating that participants actively engaged in this task. In Study 2, participants spent on average 352 trials (*SD* = 123) on the labor task; they switched 8.77 times (*SD* = 6.58) from labor to leisure. In the labor task, they responded in 84% of trials (*SD* = 6%) correctly. During the leisure task, they responded in 28% (*SD* = 17%) of trials. We conclude that in both studies, participants were engaged in both tasks, even though performance on the leisure task had no impact on their payout. See S3 for details on accuracy directly before switches to leisure, S4 for differences in accuracy before compared to after periods of leisure, and S5 for correlations of overall baseline pupil diameter and pupil dilations with performance measures.

#### Overall baseline and dilation differences between tasks

To ensure that changes in pupil baselines or dilations around switches between tasks were not due to overall differences between tasks, we fitted LMEMs to the maximum-normalized baseline and dilation scores with *task* as the single predictor (including full random effects structures). In both studies, baselines (Study 1: M_labor_ = 0.58, M_leisure_ = 0.54, Study 2: M_labor_ = 0.57, M_leisure_ = 0.56) and dilations (Study 1: M_labor_ = 0.36, M_leisure_ = 0.35; Study 2: M_labor_ = 0.36, M_leisure_ = 0.32) tended to be higher during labor compared to leisure. However, these differences between tasks were only significant for dilations in Study 2, *F*(1, 27.51) = 4.50, *p* = .043 (all other *p*s > .11), and never exceeded ∼ 0.04 on a maximum-normalized scale. These differences are substantially smaller than baseline increases and dilation decreases around switches (see below), and thus cannot explain the observed pupil changes around switches. Furthermore, if there were no additional mechanisms driving pupil changes around switches, the finding of overall slightly higher baselines and dilations during the labor task would make us expect decreases in both measures around labor-to-leisure switches, but increases in both measures around leisure-to-labor switches, which is inconsistent with our findings reported below.

### Pre-registered confirmatory analyses of Study 1

In our original pre-registration, we decided to select the last ten but one trials before labor-to-leisure switches, and fit LMEMs with the outcomes baseline pupil diameter or pupil dilations and the sole predictor trial number relative to the switch (−10 till −2). There was no evidence for an increase in baseline pupil diameter, β = .01, 95% CI [−0.04, 0.06], *F*(1, 23.50) = 0.11, *p* = .743, nor for a decrease in pupil dilations, β = -.03, 95% CI [−0.08, 0.02], *F*(1, 25.68) = 1.33, *p* = .260, during the pre-registered time window.

### Exploratory analyses Study 1 and pre-registered analyses Study 2

As a next step, we ran exploratory analyses on the data from Study 1. For this purpose, we (a) considered a different range of trials, namely the last five trials before and the first five trials after switches, centered on the trial on which participants decided to switch. This allowed us to investigate whether pupil changes started later than we initially expected, namely only 1-2 trials before the switch. Also, we (b) investigated both switches from labor to leisure and from leisure to labor. Unless otherwise indicated, all analyses reported below were pre-registered for Study 2.

#### Changes in baseline diameters

Table 1 reports the results of the fitted GAMMs and LMEMs for both studies. GAMMs yielded significant changes in both baseline pupil diameter and pupil dilations, for both switch types, in both studies. Time courses of the GAMMs are plotted in Figure 1. In Table 1, we further report LMEMs. We used these LMEMs to test whether there was a significant net increase or decrease (recognizable by the sign of the beta coefficient) across the entire specified trial window.

**Table 1.** Results of generalized additive mixed models (GAMMs) and linear mixed-effects models (LMEMs) for pupil baseline diameter and pupil dilations across labor-to-leisure and leisure-to-labor switches in Study 1 and 2.

| Measure | Switch Type | Study | GAMM | LMEM |
| --- | --- | --- | --- | --- |
| Baseline | Labor-to-leisure | 1 | $F(8.54, 3577.26) = 23.65, p < .001$ | $\beta = .12, 95\% \text{ CI } [.04, .19], F(1, 25.88) = 9.89, p = .004$ |
| | | 2 | $F(8.30, 4257.04) = 11.43, p < .001$ | $\beta = .03, 95\% \text{ CI } [-.02, .08], F(1, 26.14) = 1.56, p = .222^a$ |
| | Leisure-to-labor | 1 | $F(8.50, 3577.26) = 23.95, p < .001$ | $\beta = .16, 95\% \text{ CI } [.09, .24], F(1, 27.85) = 19.35, p < .001$ |
| | | 2 | $F(8.57, 4257.04) = 21.46, p < .001$ | $\beta = .12, 95\% \text{ CI } [.07, .17], F(1, 23.46) = 23.46, p < .001$ |
| | Difference | 1 | $F(1.00, 3584.57) = 1.16, p = .281$ | $\beta = -.02, 95\% \text{ CI } [-.08, .03], F(1, 26.56) = 0.96, p = .336$ |
| | | 2 | $F(3.33, 4261.67) = 4.06, p = .005$ | $\beta = .04, 95\% \text{ CI } [-.09, .01], F(1, 26.72) = 3.51, p = .072$ |
| Dilation | Labor-to-leisure | 1 | $F(7.76, 4012.88) = 2.91, p = .003$ | $\beta = -.07, 95\% \text{ CI } [-.14, -.01], F(1, 27.75) = 4.90, p = .035$ |
| | | 2 | $F(1, 4688.12) = 13.31, p < .001$ | $\beta = -.06, 95\% \text{ CI } [-.02, .11], F(1, 26.43) = 6.71, p = .015,$ |
| | Leisure-to-labor | 1 | $F(7.91, 4012.88) = 5.13, p < .001$ | $\beta = .001, 95\% \text{ CI } [-.07, .06], F(1, 28.17) = 0, p = .962^a$ |
| | | 2 | $F(7.66, 4688.12) = 5.24, p < .001$ | $\beta = .01, 95\% \text{ CI } [-.02, .05], F(1, 20.07) = 0.51, p = .482^a$ |
| | Difference | 1 | $F(1.80, 4002.19) = 4.16, p = .019$ | $\beta = -.04, 95\% \text{ CI } [-.09, .02], F(1, 28.54) = 1.95, p = .174^b$ |
| | | 2 | $F(7.76, 4687.73) = 6.29, p < .001$ | $\beta = -.03, 95\% \text{ CI } [-.06, -.01], F(1, 520.93) = 7.79, p = .005^b$ |

*Note.* We fitted separate models for labor-to-leisure and leisure-to-labor switches testing our pre-registered hypotheses, and then exploratory models to compare both switch types.
<sup>a</sup> Estimates were significantly different from zero when we re-fitted them on a more constrained time window of two trials before until two trials after switches (see main text).
<sup>b</sup> Due to convergence warnings, we simplified the model by dropping the random slopes either per bout or per subject until the model converged.

As predicted by both adaptive gain theory and network reset theory, GAMMs showed that baseline pupil diameter increased around labor-to-leisure switches in both Study 1 and Study 2 (Figures 2a and b). Whereas LMEMs indicated that there was also an overall net increase in baselines in Study 1, there was no significant evidence for such an increase in Study 2. Nevertheless, Figure 2b clearly displays an increase, which might be too temporally constrained to be picked up as a net increase by the LMEM. We thus ran an additional exploratory analysis by refitting this LMEM on a more restricted time window from two trials before until two trials after the switch, which yielded indeed a strong, but temporally constrained, increase, β = .13, 95% CI [.07, .20], *F*(1, 27.30) = 13.19, *p* = .001.

**Figure 2.**
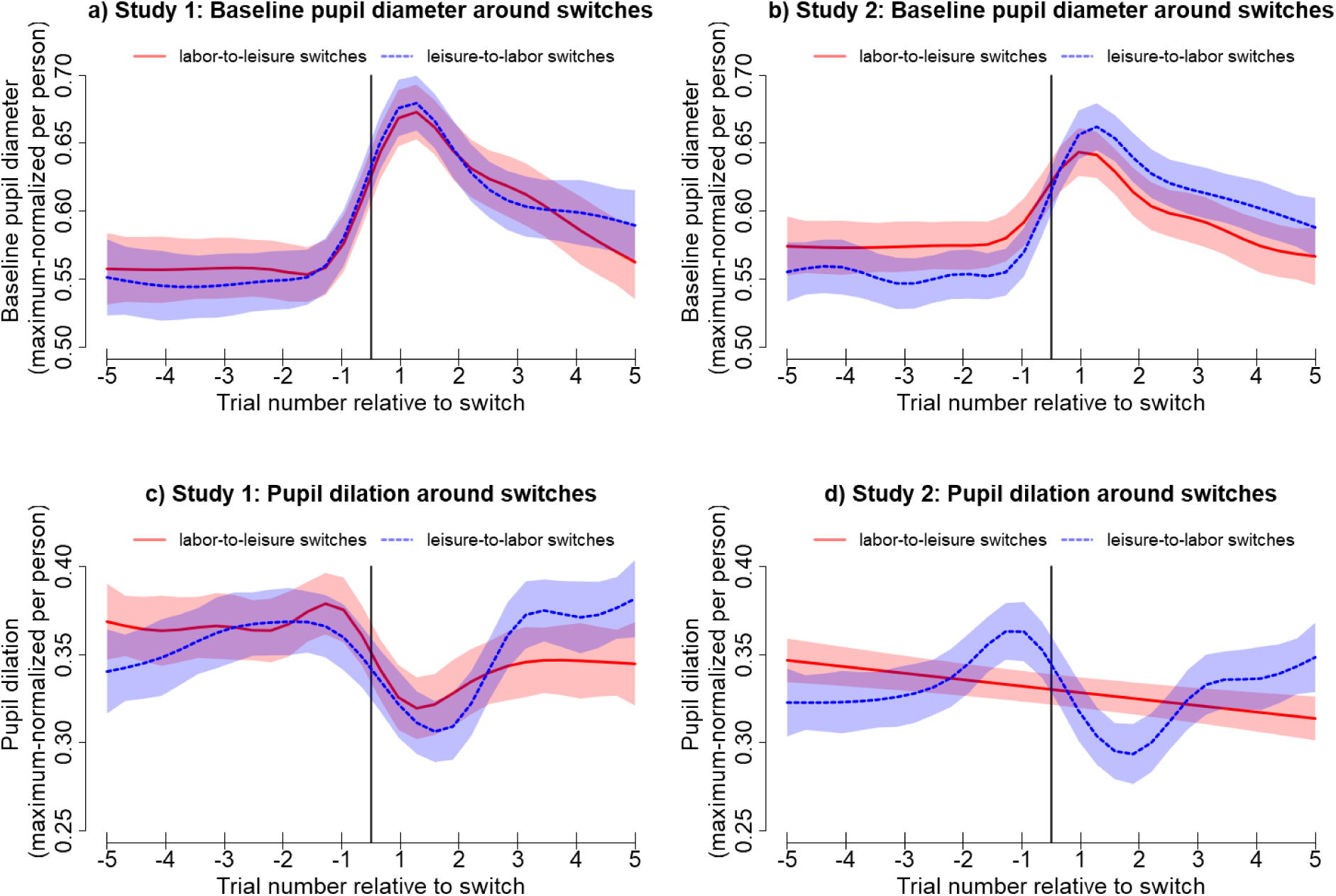
Top panels: Time course of baseline pupil diameter around labor-to-leisure and leisure-to-labor switches in Studies 1 (A) and 2 (B). Bottom panels: Time course of pupil dilations around labor-to-leisure and leisure-to-labor switches in Studies 1 (C) and 2 (D). All time courses were derived from the GAMMs described in Table 1 (time courses are model estimates, not averages of data). Shades indicate 95%-CIs. Vertical lines indicate the time point of the switch.

**Figure 3.**
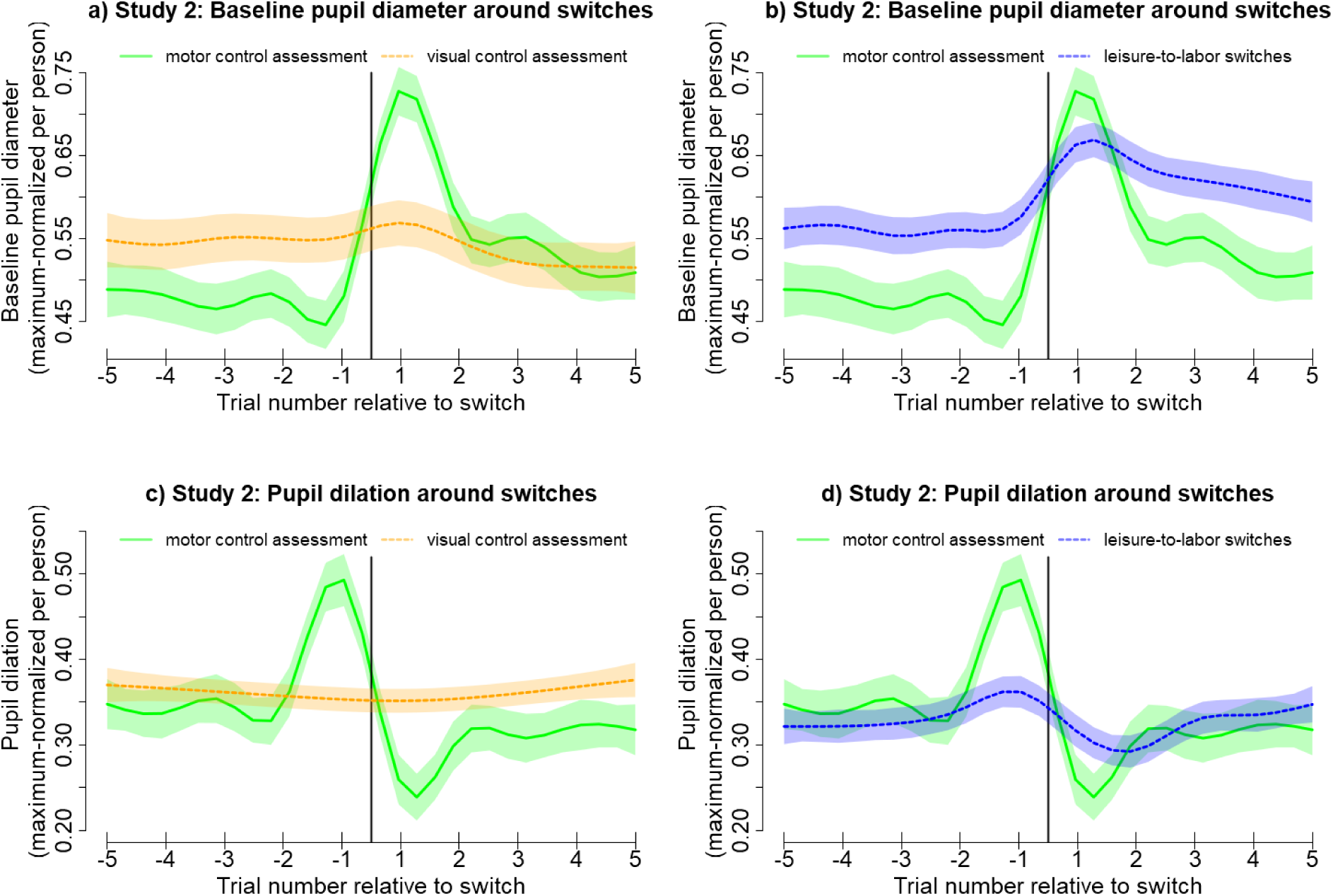
Time course of in pupil baseline diameter (a) and pupil dilations (c) in the motor and visual control assessment in Study 2. For both baselines and dilations, there was a significant increase and subsequent decrease around switches in the motor condition, but not in the visual condition. The baseline changes in the motor condition were more peaked, but also more short-lived than those in leisure-to-labor switches (b). The dilation increase in the motor condition was unparalleled in leisure-to-labor switches (d). Plots are based on the GAMMs computed. Shades indicate 95%-CIs. Vertical lines indicate the time point of the switch.

Furthermore, in line with network reset theory, but in contradiction with adaptive gain theory, GAMMs yielded a similar increase in baselines across leisure-to-labor switches in both studies. This was corroborated by LMEMs in both studies, which both showed net increases in baselines across these switches.

We ran exploratory analyses to test whether the time courses of pupil dilations differed between switch types. When comparing switch types in Study 1, neither GAMMs nor LMEMs indicated evidence for a significant difference between switch types. In Study 2, the GAMM yielded that the increase in baselines was even slightly stronger for leisure-to-labor compared to labor-to-leisure switches (see Figure 2b), while the LMEM yielded no evidence for a net difference. Overall, in both studies, we found significant increases in pupil baselines in labor-to-leisure switches, which is consistent with both adaptive gain theory and network reset theory, as well as in leisure-to-labor switches, which is consistent with network reset theory, but inconsistent with adaptive gain theory.

#### Changes in pupil dilations

For labor-to-leisure switches, as predicted by both adaptive gain theory and network reset theory, GAMMs revealed significant decreases in pupil dilations in both studies (see Figures 2c and d). This was corroborated by LMEMs indicating significant net decreases in dilations in both studies.

For leisure-to-labor switches, GAMMs indicated a similar decrease in both studies, which was in line with network reset theory, but inconsistent with adaptive gain theory. However, LMEMs did not yield evidence for a significant net decrease in either study. Given that decreases were clearly visible in Figures 2c and d, we again ran exploratory analyses by refitting the models on a more temporally restricted trial window from two trials before until two trials after a switch. In this window, there was a strong significant decrease both in Study 1, β = -.16, 95% CI [−.24, -.08], *F*(1, 21.94) =15.36, *p* < .001, and Study 2, β = -.13, 95% CI [−19, -.07], *F*(1, 21.07) = 17.41, *p* < .001.

We ran exploratory analyses to test whether the time courses of pupil dilations differed between switch types. As can be seen in Figure 2c, GAMMs indicated that dilation time courses were significantly different between switch types in Study 1, with dilations higher three to five trials after leisure-to-labor switches compared to labor-to-leisure switches. The respective LMEM yielded no evidence for net differences. In Study 2, both GAMMs and LMEMs indicated significant differences between switch types: Figures 2d shows that dilations were higher five trials before and two trials after labor-to-leisure compared to leisure-to-labor switches, but lower one trial before and five trials after labor-to-leisure compared to leisure-to-labor switches.

We make two more observations about the particular shape of the pupil dilation time courses across switches. First, note that in in both studies, before the decrease, there seems to be a brief increase on the last trial before a switch (see Figures 2c and d). Although neither adaptive gain theory nor network reset theory predicts this increase, it may be explained by the idea that pupil dilation reflects effort mobilization (Beatty & Lucero-Wagoner, 2000; Hess & Polt, 1964; Kahneman & Beatty, 1966; Laeng, Sirois, & Gredebäck, 2012): it may be the case that initiating the task switch requires particular effort (see also the motor control assessments below).

Second, regarding labor-to-leisure switches, the GAMM for Study 1 displays a somewhat abrupt decrease that is constrained to one to two trials around switches (Figure 2c, red line), while the GAMM for Study 2 (Figure 2d, red line) displays a more gradual decrease over the entire trial window. Here again, neither adaptive gain theory nor network reset theory make predictions about the exact shape of the decrease. However, the difference between studies may be explained from the way GAMM models fit smooth curves through data. In GAMM models, the order of the best-fitting polynomial is determined by trading off model fit vs. model complexity. For a noisier raw data pattern (with multiple peaks and troughs), this trade-off may favor a lower order polynomial (as in Figure 2d), which is more likely to reflect the data generative process. When we visually inspected pupil dilation time courses (for the script generating the respective plots, see https://osf.io/ukgsh/), we did indeed observe that the dilation data for labor-to-leisure switches seemed to be noisier in Study 2 than Study 1, which may explain why the decrease in pupil dilation seemed to be more gradual in Study 2.

#### Pupil changes around switches compared to overall differences between tasks

A possible explanation for changes in pupil baselines and dilations around task switches is that those measures are overall higher in one task compared to the other. If this was the case, pupil changes around switches would merely reflect the transition from one task to the other, rather than particular processes that are involved in implementing the task switch. To rule out this possibility, we compared the magnitude of pupil changes around switches (see Figure 2) to overall differences between tasks on those measures. Baseline increases around switches were of magnitudes ∼0.15 on the maximum-normalized scale and dilation decreases of magnitudes ∼0.10, which was both substantially larger than overall differences between tasks on those measures (which did not exceed ∼0.04). We thus deem it unlikely that the observed baseline and dilation changes around switches are reducible to overall differences between tasks on those pupil measures.

#### Summary of main results

In conclusion, across both studies, we found increases in pupil baseline levels and decreases in pupil dilations around both labor-to-leisure and leisure-to-labor task switches. The size of baseline increases and dilation decreases was largely comparable between both switch types and across both studies.

These results were fully predicted by network reset theory. Only the patterns observed for labor-to-leisure switches, but not the pattern for leisure-to-labor switches was consistent with adaptive gain theory. In sum, our data provided support for network reset theory, which proposes that pupil shifts across both types of task switches reflect processes involved in task-switching, independent of the motivational nature of the task. However, one might argue that these shifts are comparable across switches types and studies because they do not reflect neural processes underlying task switching, but rather a) motor processes initiating the movement of the joystick, or b) visual processes of observing the frame highlighting the current task moving from one task label to the other. Although these alternative explanations seem rather unlikely given the prolonged elevation of pupil baselines until twelve seconds after the behavioral switch, we aimed to rule out those alternative explanations via two additional assessments.

### Control assessments (Study 2 only)

#### Motor control assessment

A GAMM indicated that contrary to our expectations, the time course of baseline pupil diameter around motor actions was significantly different from zero, *F*(8.93, 1535.64) = 40.40, *p* < .001 (see Figure S6a), and a LMEM yielded a significant increase, β = .11, 95% CI [.05, .16], *F*(1, 22.00) = 15.25, *p* < .001. Additionally, contrary to our expectations, a GAMM indicated that the time course of pupil dilation was significant different from zero, *F*(8.94, 1665.19) = 20.47, *p* < .001 (see Figure S6c), with a LMEM yielding a significant decrease, β = -.08, 95% CI [−.12, -.03], *F*(1, 21.99) = 10.63, *p* = .004.

Several factors might account for this unexpected finding. First, we expected this condition to be an easy task, but participants committed a considerable amount of errors (217 out of 2,480 trials, i.e. 10% of all trials; with 148 false-positive switches and 69 false-negative non-switches).^5^ This indicates that the task was more difficult (or less engaging) than expected, and effortful processes different from mere motor activation (e.g., working memory) might have contributed to pupil changes (Kahneman, 1973). Second, it seemed that pupil dilations were particularly high in the motor control assessments on the very last trial before a switch, i.e. the trial participants saw the face that triggered them to switch, which could reflect increased effort (Hess & Polt, 1964; Kahneman & Beatty, 1966). In an exploratory analysis, we directly contrasted the time courses in the motion control assessment with those around leisure-to-labor switches,^6^ using a GAMM with the interaction between trial number and the ordered factor switch type (motor control assessment vs. leisure-to-labor). The differences in the time course of pupil dilation was indeed significant, *F*(8.93, 3975.57) = 12.88, *p* < .001, indicating that pupil dilations were higher in the motor control assessment on the very last trial before a switch, but higher in leisure-to-labor switches on the very first trial after a switch (see Figure S6d). Third, it seemed that baseline increases in motor control assessments were restricted to the very first trial after a switch, but more sustained in labor-to-leisure and leisure-to-labor switches. When directly contrasting the time courses of pupil baselines of motor control assessments and leisure-to-labor switches, we indeed found significant differences, *F*(8.57, 3657.29) =12.27, *p* < .001: baselines were higher in leisure-to-labor switches from five till one trial before the switch and from two to five trials after the switch, but the motor control assessment was only higher on the very first trial after a switch (see Figure S6b).

Taken together, the motor control assessment might have been more difficult for participants than we expected, as indicated by the high error rates and the strong pupil dilation on the last trial before the switch, which might reflect increased effort recruitment. Also, increases in the labor-to-leisure and leisure-to-labor switches continued for several trials after a switch, while the increase in the motor control assessment was only present on the very first trial after a switch. These findings render it unlikely that the pupil changes observed in labor-to-leisure and leisure-to-labor switches were fully reducible to processes underlying motor movements.

### Visual control assessment

In the selected time window, contrary to our expectations, a GAMM indicated that the time course of baselines in the visual control assessment was significantly different from zero, *F*(6.63, 2075.75) = 2.92, *p* = .006. Visual inspection showed however that the time course was overall rather flat (Figure S6a).^7^ A LMEM found no evidence for any net changes across the selected time window, β = -.04, 95% CI [−.09, .01], *F*(1, 30.00) = 3.63, *p* = .067. Similarly, a GAMM found no evidence for the time course of pupil dilations deviating from a flat line, *F*(2.48, 2335.32) = 2.09, *p* = .112 (see Figure S6b), and a LMEM found no evidence for net changes either, β = .01, 95% CI [−.03, .04], *F*(1, 30.00) = 0.04, *p* =.835. We conclude that the pupil changes observed in labor-to-leisure and leisure-to-labor switches do not stem from merely watching the frame highlighting task labels move.

### Correlations with action orientation across both studies

We initially pre-registered to run LMEMs with the predictors trial number, switch type, the respective questionnaire scale, and all interactions to check whether individual differences in procrastination and action orientation predicted the amplitude of pupil changes, and differently so for the different switch types. Since these models failed to converge, we instead re-used the simple models of baseline increases and dilation decreases reported above, extracted the best linear unbiased predictors (BLUPs),^8^ i.e. the group-level fixed effect slope of trial number plus the respective random slope of trial number of each participant, and correlated those with participants’ average scores on the questionnaires. Note that we did not compare correlations against zero, but against the correlations of the respective other switch type. Correlations between BLUPs and questionnaires are displayed in Table 2; descriptive statistics and intercorrelations between questionnaires in SOM S7. Note that pupil dilation decreases around switches, so BLUPs are typically negative; the more negative they are, the larger is the amplitude of the decrease. Most correlations were low. As the most noteworthy finding, which we think might warrant follow-up research, we observed that the BLUPs of pupil dilation decreases were positively associated with action orientation after failure (AOF) and self-regulation (SR) for labor-to-leisure switches, but negatively for leisure-to-labor switches. The same pattern, though weaker, was found for the subscales decision-related action orientation (AOD) and self-control (SC).^9^ The negative correlations with BLUPs in leisure-to-labor switches imply that the higher participants scored on action orientation, the stronger were their decreases in pupil dilation in leisure-to-labor switches, but the weaker were their decreases in pupil dilation in labor-to-leisure switches. However, given our limited sample size of only 62 participants, those findings might not be robust and need further corroboration by future research. No associations were found with procrastination (IPS) and volitional development (VD).

**Table 2.** Correlations of Self-Reported Procrastination Tendencies and Action vs. State Orientation With the Best Linear Unbiased Predictors (BLUPs) of Pupil Baseline Diameter Increases and Pupil Dilation Decreases in Both Labor-to-Leisure and Leisure-to-Labor Switches

|  | BLUPs Baseline |  | BLUPs Dilation |  |
| --- | --- | --- | --- | --- |
|  | Labor-to-leisure switches | Leisure-to-labor switches | Labor-to-leisure switches | Leisure-to-labor switches |
| IPS | -.04 | .01 | -.14 | .01 |
| ACS-24 AOF | .17 | -.09 | .19 | -.28 |
| ACS-24 AOD | .14 | .05 | .07 | -.13 |
| SSI K3 SR | .07 | -.11 | .26 | -.27 |
| SSI K3 SC | -.04 | .20 | .16 | -.25 |

<sup>9</sup> All these patterns were also reflected in significant or marginally significant 3-way interactions in the initially pre-registered LMEMs, which however yielded convergence warnings.

|  |  |  |  |  |
| --- | --- | --- | --- | --- |
| SSI K3 VD | -.01 | -.14 | .10 | .03 |

*Note.* Higher IPS values indicate higher procrastination tendencies, and higher values on the other scales indicate higher action orientation. Samples of both studies (total $N = 62$ ) were combined. IPS = Irrational Procrastination Scale; ACS-24 AOF = ACS-24 Action orientation subsequent to failure vs. preoccupation subscale; ACS-24 AOD = ACS-24 Prospective and decision-related action orientation vs. hesitation subscale; SSI-K3 SR = SSI-K3 Self-regulation (Competence) subscale; SSI-K3 SC = SSI-K3 Self-control subscale; SSI-K3 VD = SSI-K3 Volitional development (Action development) subscale.
† $p < .10$ , \* $p < .05$ , uncorrected.

## Discussion

In exploratory analyses (Study 1) and confirmatory analyses (Study 2), we found that pupil baseline levels increased and pupil dilations decreased around switches from cognitive labor to cognitive leisure, and from leisure to labor. These changes in pupil baseline levels and pupil dilations were short-lived. Our findings extend previous studies that found similar pupil changes when people disengaged from (and restarted) an auditory discrimination task (Gilzenrat et al., 2010), when people shifted between multiple bandit gambling machines (Jepma & Nieuwenhuis, 2011) and when people shifted between strategies in solving Raven’s Matrices (Hayes & Petrov, 2016). By contrast to these previous studies, in our research, participants chose between two motivationally different tasks—namely an effortful, but profitable 2-back (labor) and an effortless, but unprofitable attractiveness rating task (leisure). Participants showed the same pupillary changes around switches between tasks in either direction. We will now discuss the main findings in greater detail.

### Pupil changes were similar for labor-to-leisure and leisure-to-labor switches

Similar pupil changes (increased baseline, decreased dilations) occurred around switches from labor to leisure *and* in switches from leisure back to labor. While the former finding (labor to leisure) is consistent with adaptive gain theory, the latter finding (leisure to labor) is not. Thus, it seems that pupillary changes around task switches cannot be interpreted as reflecting people’s motivation to take a break. Rather, these pupillary changes might reflect cognitive processes that underlie task switches more generally. This interpretation is in line with some predictions from network reset theory, which suggests that increases in pupil size (putatively reflecting NE release) occur when reorientation towards a new environment is needed. This reorientation account fits previous research that found increases in pupil diameter in response to environmental instabilities and surprises (Lavín, San Martín, & Rosales Jubal, 2014; Nassar et al., 2012; Preuschoff, ’t Hart, & Einhäuser, 2011) and in response to rule changes in the Wisconsin Card Sorting Task (Pajkossy, Szőllősi, Demeter, & Racsmány, 2017). Indeed, recent research found increased BOLD signals in the LC during task switching in humans (von der Gablentz, Tempelmann, Münte, & Heldmann, 2015).

Thus, at least on first sight, our findings seem more consistent with predictions from network reset theory, than with predictions from adaptive gain theory. We hasten to add, however, that it is debatable whether our data are in line with all aspects of network reset theory. In particular, network reset theory is typically used to understand how unexpected changes in environmental requirements drive NE-related reorientation of cortical networks (and its downstream behavioral consequences). It is not yet clear to what extent network reset theory can help to model the effects of people’s self-initiated decisions, too.

Nevertheless, research does lean towards the idea that network reset theory applies to self-initiated decisions. In particular, one author of network reset theory (Sara, 2015, 2016) recently cited evidence for LC involvement in a task switching paradigm in humans (von der Gablentz et al., 2015) as evidence for network reset theory. In this paradigm, participants were not explicitly informed about task switching requirements. Instead, they had to infer when they needed to switch from performance feedback and from their own confidence in having responded correctly. Thus, it appears that also task switches that do not follow from evident changes in the environment, but from an inference process, may activate the LC. Using the same rationale, our results might extend network reset theory by suggesting that NE responses to the detection of environmental changes, such as threats or dangers, might also occur as responses to people’s self-initiated decision to switch between tasks, perhaps helping them to adapt to the new task’s requirements.

Related to the latter line of reasoning, some recent studies suggest that NE plays a more active role in updating action policies (O’Reilly et al., 2013; Urai, Braun, & Donner, 2017; Van Slooten, Jahfari, Knapen, & Theeuwes, 2018), helping people to adopt new task mindsets, rather than (or in addition to) the role of encoding environmental uncertainty (Dayan & Yu, 2006; Lavín et al., 2014; Nassar et al., 2012). Such a proposed role for NE in reconsidering action policies is consistent with recent electrophysiological work in monkeys that showed that (a) the activity of dopaminergic neurons in the substantia nigra correlates with motivational variables such as expected levels of reward and effort, potentially integrating different signals into one single decision, but (b) activity of NE neurons in the LC correlates with actual effort exertion, potentially aiding the execution of this decision (Varazzani, San-Galli, Gilardeau, & Bouret, 2015).

In sum, building on our findings and on recent insights in NE function, it is possible that previous experiments about shifts from exploitation to exploration (Gilzenrat et al., 2010; Hayes & Petrov, 2016; Jepma & Nieuwenhuis, 2011; Kane et al., 2017; Pajkossy et al., 2017) observed NE facilitating the execution of task switching, rather than NE facilitating motivational shifts. By contrast to these previous studies, our paradigm used two qualitatively distinct tasks, with distinct motivational properties (i.e., labor vs. leisure). Due to this design feature, our study could reveal that the motivational direction does not matter.

### Pupil changes were short-lived

Our studies provide a fine-grained examination of the time course of baseline pupil diameter and pupil dilations around task switches. For both switches from labor to leisure and vice versa, changes in baselines and dilations began later than initially expected. In particular, baselines started to increase on the last trial (3 seconds) before a switch and levelled off around four trials (12 seconds) after the switch (see Figure 2a and b). Generally, dilations decreased from one trial before (3 seconds) until two trials after a switch (6 seconds) and then returned to pre-switch levels (exception: labor-to-leisure switches did not show this effect in Study 2, potentially due to measurement noise). Overall, changes in baselines and dilations were comparable between both switch types and across both studies. Importantly, both baselines and dilations returned to pre-switch levels after a few trials, which was reflected in that our linear mixed-effect models did not always yielded a net change over the entire trial window (30 seconds).

The short-livedness of pupil changes is potentially relevant to our evaluation of adaptive gain theory and network reset theory. In particular, adaptive gain theory predicts that for extended periods of exploration, pupil baselines are permanently high and dilations permanently low. Such extended periods of exploration may take the form of *mind wandering*, during which people may plan for future activities (see Kool & Botvinick, 2014). In line with this idea, some studies found increased pupil baselines during mind-wandering (Franklin, Broadway, Mrazek, Smallwood, & Schooler, 2013), while others found lower baselines (e.g., Grandchamp, Braboszcz, & Delorme, 2014; Konishi, Brown, Battaglini, & Smallwood, 2017; Mittner et al., 2014; Unsworth & Robison, 2016, 2018). Network reset theory, by contrast, predicts rapid adaptations around task switches. In this regard, our data seem to consistent with predictions derived from network reset theory, as well.

### Discoveries from exploratory analyses

Exploratory analyses suggested that individual differences in pupil dilation changes around switches were correlated with individual differences in self-reported action orientation. In particular, participants with higher action orientation showed smaller decreases in pupil dilation around switches from labor to leisure, but larger decreases in pupil dilation around switches from leisure to labor. This association might be plausible when the neural processes reflected by pupil dilation decreases help humans inhibit an old task mindset and prepare a new one, as proposed by network reset theory. In this case, inhibition and preparation are especially needed when going back to labor, as labor is both effortful and returns rewards, but not when switching to leisure. One might speculate that people who more selectively use their reorientation mechanisms in situations in which they are actually needed, i.e. when preparing an effortful task, may also approach challenges more proactively in everyday life.

In line with the latter line of reasoning, previous research indeed found action-oriented individuals to use their working memory capacity more effectively (Jostmann & Koole, 2007) than state-oriented individuals. A similar association was recently discussed for fluid intelligence (Hayes & Petrov, 2016), suggesting that more intelligent individuals show stronger pupil dilations compared to average-intelligent individuals in challenging tasks (van der Meer et al., 2010), but weaker dilations in simple tasks (Ahern & Beatty, 1979). However, given that we did not predict this association a-priori, these results are should be considered exploratory, and our interpretation should be considered speculative, requiring further corroboration by future work. Worth noting, no such correlations were found for differences in baseline increases. Statistical power for these correlational analyses was quite low in our study. Thus, future research needs to clarify the role of individual differences, with individual differences in action orientation as a potential starting point.

### Conclusions and future directions

In sum, our results show that the pupil dynamics that are often thought to reflect shifts from exploitation to exploration, also occur in shifts from exploration back to exploitation. Our results also indicated that these pupillary changes were short-lived; they were disappeared within seconds. Though with caution, we suggest that pupillary shifts around task switches may not reflect a motivational process (people wanting to take a break), but a reorientation process (people preparing for the new task).

A limitation of our approach is that we used research on the exploitation-exploration trade-off to generate hypotheses about the labor-leisure trade-off. As noted in the introduction, those two trade-offs might co-occur under particular circumstances, but are prima facie independent. While the mapping of our tasks on the categories labor and leisure is likely stable, the mapping on exploitation and exploration might vary over time, e.g. when during the leisure task, participants suddenly detect the chance to indulge in a more abstract, intrinsic reward, such as a pleasant memory of an event outside the task context. Thus, more theoretical work—potentially using computational modeling to estimate the dynamic values of different goals, including extrinsic, monetary goals and intrinsic, more hedonic goals—is needed to draw conclusions about when a certain activity constitutes an exploitative option and when not (Meyniel et al., 2013; Mittner et al., 2014).

Future research might more directly test whether changes in pupil baselines and dilations are independent from the motivational properties of tasks. Also, future research might include control experiments that test whether the same pupil changes occur when task switches are externally induced (e.g., when participants see a countdown that requires them to switch). Such a study could elucidate whether externally-induced switches (as classically concerned by network reset theory) and self-directed switches as implemented in our task share common mechanisms, e.g., the updating of action policies. Finally, it might be interesting to further investigate whether the magnitude of pupil changes is moderated by the difficulty of an upcoming task or by individual differences in action orientation.

## Data and Code Availability

Materials used in data collection as well as data and code to reproduce our analyses are available under https://osf.io/b9z4c/ (Study 1) and https://osf.io/ukgsh/ (Study 2).

## Acknowledgments

The authors would like to thank H. Voogd for assistance with programming the eye-tracker and J. Kuhl for his kind permission to use the questionnaires he created. J.A. would like to thank the Behavioural Science Institute at Radboud University for funding the research of his Master’s thesis. E. B. was supported by grant 016-165-100 from the Netherlands Organization for Scientific Research.

## Supplementary Online Materials

## Appendix S1

### Participant- and Trial-Based Data Inclusion in Study 1

Four of the 35 participants were excluded from all analyses in accordance with our pre-registered criteria: Two of them did never switch to the leisure task, but only performed the labor task; one performed below 50% accuracy in the labor task, and one participant’s eye-tracking data were unusable due to a high percentage of missing values.

For the *pre-registered* analyses of pupil data, we selected the last ten but one trials before each labor-to-leisure switch. The adjacent trials preceding the same switch were treated as belonging to the same *bout*, with trials nested in bouts and bouts nested in participants, yielding 201 bouts (1,809 trials).

For the *exploratory* analyses of pupil data, we selected the last five trials before each switch and the first five trials after each switch. The trials around one single switch were treated as belonging to the same *bout* of adjacent trials, with trials nested in bouts and bouts nested in participants. This resulted in a final data set with 247 labor-to-leisure switches (2,225 trials) and 247 leisure-to-labor switches (2,180 trials).

## Appendix S2

### Participant- and Trial-Based Data Inclusion in Study 2

In line with our pre-registered criteria, we excluded two participants from all our analyses as they only performed the labor task and never switched, and two further participants because their eye-tracking data were unusable due to high rates of missing values. This left a sample of 31 participants that was used in all analyses.

We selected again the last five trials before each switch between tasks and the first five trials after each switch. Here, also the joystick movements in the motor control assessment phase and the observed frame movements in the visual control assessment were treated as switches. In total, we analyzed 272 switches from labor to leisure (2,594 trials), 273 switches from leisure to labor (2,535 trials), and 248 visual switches (2,480 trials).

Regarding the motor control assessment, unexpectedly, participants committed a considerable amount of errors (217 out of 2,480 trials, i.e. 10% of all trials; of which 148 were false-positive switches and 69 false-negative non-switches). As a liberal exclusion criterion, to keep as many correct trials as possible, we decided to first exclude eight participants, as each of their eight bouts, i.e. required movements of the joystick surrounded by ten trials, contained at least one error. This left 39 trials with errors (11 false positives, 28 false negatives), which we deleted in a second step, resulting in 164 remaining bouts (1,880 trials). As a more conservative exclusion criterion, we excluded all bouts with any errors, resulting in 155 bouts (1,550 trials). The pattern of significant and non-significant results was the same for both exclusion criteria. In the main text, we report results based on the liberal exclusion.

## Appendix S3

### Testing Accuracy Decrements Before Breaks

A potential reason for participants’ decision to switch from the labor to the leisure task might be that they recognize a decrease in their performance, are afraid that they might forfeit their chance for an extra monetary bonus, and thus decide to take a break in order to rest and restore their ability to concentrate. Under this explanation, one might assume that switches could be induced by error monitoring, independent of changes in motivation or NE levels.

However, note that participants did not receive any error feedback during the tasks, and thus had imperfect knowledge about their performance. Furthermore, AGT itself predicts that increased tonic NE levels will lead to decreases in accuracy, which is supported by behavioral evidence: animal work found switches from exploitation to exploration to be accompanied by increased false alarm rates (Cohen, McClure, & Yu, 2007), and increased pupil diameter has been found associated with lapses in attention in humans (Hopstaken, van der Linden, Bakker, & Kompier, 2015; Unsworth & Robison, 2016). Hence, decreases in accuracy might not be an independent mechanism, but a side-effect of changes in NE levels.

In exploratory analyses, we tested whether decreases in accuracy would occur during the last five trials before switches from labor to leisure. We used generalized linear mixed effects models (GLMEMs) in the package lme4 (Bates, Mächler, Bolker, & Walker, 2015) in R (R Core Team, 2017) with accuracy (binary: correct or incorrect) as outcome and trial number relative to switch as predictor. The predictor was standardized. We added a random intercept and a random slope of trial number for each participant and for each bout of each participant to the model, including all possible random correlations. *P*-values were calculated using likelihood ratio tests as implemented in the package afex (Singmann, Bolker, Westfall, & Aust, 2018), since *F*-tests are not available for GLMEMs. We computed 95% confidence intervals via bootstrapping with 1,000 simulations using lme4’s bootMer function, which uses the boot package (version 1.3.18, Canto & Ripley, 2016). Effectively, we analyzed of 376 switches (1,525 trials) in Study 1 and 372 switches (1,591 trials) in Study 2.

In Study 1, there was a non-significant trend of decreasing accuracy during the last five trials before switches from labor to leisure, β = -.10, 95% CI [−.25, .05], χ^2^(1) = 2.85, *p* =.091. In Study 2, there was a significant decrease in accuracy, β = -.26, 95% CI [−.47, -.06], *χ^2^*(1) = 4.90, *p* = .027. We conclude that there was mixed evidence for accuracy decreasing directly before a break.

## Appendix S4

### Testing Accuracy Restoration After Compared to Before Breaks

As explained in S3, subjects might use breaks as a chance to restore their ability to concentrate. We thus tested in exploratory analyses whether accuracy was significantly better in the first five trials after a break (i.e. a series of leisure trials) compared to the last five trials before a break.

We used GLMEMs with accuracy as outcome and a binary factor indicating whether trials were before or after the leisure bout as the sole predictor. Treatment-coding was applied. We added random intercepts and slopes of trial number for each participant to the model, including all random correlations. Again, *p* values were computed with likelihood ratio tests. We analyzed 247 breaks (2,181 trials) in Study 1 and 272 breaks (2,338 trials) in Study 2.

In Study 1, indeed, mean accuracy was slightly higher during the first five labor trials after a switch from leisure back to labor (*M* = .85, *SD* = .36), than during the last five trials before a switch from labor to leisure (*M* = .80, *SD* = .40), β = .15, 95% CI [.01, .27], χ^2^(1) = 5.23, *p* = .022. In Study 2, however, there was no significant increase in average accuracy from before a break, (*M* = .79, *SD* = .41) to after a break (*M* = .81, *SD* = .39), β = .03, 95% CI [−.11, .16], *χ^2^*(1) = 0.63, *p* = .427. Hence, we obtained mixed evidence for the hypothesis that performing the leisure task had a recreational function and immediately improved participants performance.

## Appendix S5

### Correlations of Overall Average Baseline Pupil Diameter and Pupil Dilations With Performance Measures Across Both Studies

| Variable | 2 | 3 | 4 | 5 | 6 |
| --- | --- | --- | --- | --- | --- |
| 1. Average pupil baselines | -.37** | -.03 | .29* | -.12 | .26* |
| 2. Average pupil dilations | - | .01 | -.19 | .01 | .08 |
| 3. # trials spent on labor |  | - | -.33** | .35** | .27 |
| 4. # switches away from labor |  |  | - | .38** | -.09 |
| 5. % button presses during labor |  |  |  | - | .38** |
| 6. % correct responses during labor |  |  |  |  | - |

*Note.* Pearson correlations of aggregated raw baseline pupil diameter and aggregated raw pupil dilations with counts of events (trials, switches) and percentages (button presses, correct responses) during labor trials across both studies ( $N = 62$ ).
\* $p < .05$ , \*\* $p < .01$ , uncorrected.

## Appendix S6

## Appendix S7

### Group-Level Means, Standard Deviations, Cronbach’s Alpha, and Correlations Between Self-Reported Procrastination Tendency and Action- vs. State Orientation

| Measure | 2 | 3 | 4 | 5 | 6 | *M* | *SD* | $\alpha$ |
| --- | --- | --- | --- | --- | --- | --- | --- | --- |
| 1. IPS | -.07 | -.67** | -.48** | -.17 | -.76** | 3.01 | 0.62 | .86 |
| 2. ACS-24 AOF | – | .47** | .58** | .17 | .09 | 4.76 | 3.26 | .81 |
| 3. ACS-24 AOD |  | – | .50** | .20 | .59** | 5.98 | 3.28 | .80 |
| 4. SSI-K3 SR |  |  | – | .33* | .28* | 2.57 | 0.42 | .80 |
| 5. SSI-K3 SC |  |  |  | – | .19 | 2.60 | 0.43 | .74 |
| 6. SSI-K3 VD |  |  |  |  | – | 2.52 | 0.48 | .84 |

*Note.* Higher IPS values indicate higher procrastination tendencies, and higher values on the other scales indicate higher action orientation. Samples of both studies (total $N = 62$ ) were combined. IPS = Irrational Procrastination Scale; ACS-24 AOF = ACS-24 Action orientation subsequent to failure vs. preoccupation subscale; ACS-24 AOD = ACS-24 Prospective and decision-related action orientation vs. hesitation subscale; SSI-K3 SR = SSI-K3 Self-regulation (Competence) subscale; SSI-K3 SC = SSI-K3 Self-control subscale; SSI-K3 VD = SSI-K3 Volitional development (Action development) subscale.
\* $p < .05$ , \*\* $p < .01$ , uncorrected.

## Footnotes

1 Pre-registration for Study 1: https://osf.io/ypyha/register/5730e99a9ad5a102c5745a8a

2 Pre-registration for Study 2: https://osf.io/48kyw/register/5730e99a9ad5a102c5745a8a

3 Maximum-normalization was implemented by subtracting the minimum of an individual from all their values and dividing them by the person’s maximum. This transformed each individual’s values into the range from 0 to 1, reducing the effect of outliers. All analyses were repeated with the raw pupil measures as well as with these measures z-standardized per person. In both cases, the same pattern of significant and non-significant results was obtained.

4 As a control, we also fitted GAMMs with random smooths, featuring an ARIMA(1)-model based on the model with random slopes to account for auto-correlation. However, given that each bout comprised at most 10 (and often less) data points, fitting separate smooths for each bout restricted us to using 8 knots for each smooth only, further constraining the “wigglyness” of the underlying smooth we could possibly fit. Auto-correlations were hardly any further reduced. These models yielded *F*- and *p*-values that were highly similar to those of the models with random slopes and led to the same conclusions.

5 For appropriate exclusion of participants and trials from the following analyses, see SOM S2.

6 Pupil changes around leisure-to-labor switches were overall larger than those around labor-to-leisure switches and should thus be more similar to the time course in the motor control assessment, providing a more conservative test for differences. The same results were found when comparing switches in the motor control assessment to labor-to-leisure switches.

7 When fitted with random smooths on eight knots, the time course in the visual control assessment was not significantly different from a flat line, *F*(2.88, 1695.85) = 2.21*, p* = 0.060.

8 BLUPs were fitted based on the total time window of interest, even though on a group-level, there was no significant decrease in dilations across leisure-to-labor switches in either study. This does not exclude the existence of meaningful individual differences between participants.

9 All these patterns were also reflected in significant or marginally significant 3-way interactions in the initially pre-registered LMEMs, which however yielded convergence warnings.

